# Genome and transcriptome analyses reveal genes involved in the formation of fine ridges on petal epidermal cells in *Hibiscus trionum*

**DOI:** 10.1101/2023.05.23.541865

**Authors:** Shizuka Koshimizu, Sachiko Masuda, Arisa Shibata, Takayoshi Ishii, Ken Shirasu, Atsushi Hoshino, Masanori Arita

## Abstract

*Hibiscus trionum*, commonly known as the ’Flower of an Hour’, is an easily cultivated plant in the Malvaceae family. The purple base part of its petal exhibits structural color due to the fine ridges on the epidermal cell surface, and the molecular mechanism of ridge formation has been actively investigated. We performed genome sequencing of *H. trionum* using a long-read sequencing technology with transcriptome and pathway analyses to identify candidate genes for fine structure formation. The ortholog of *AtSHINE1*, which is involved in the biosynthesis of cuticular wax in *Arabidopsis thaliana*, was significantly overexpressed in the iridescent tissue. In addition, orthologs of *AtCUS2* and *AtCYP77A*, which contribute to cutin synthesis, were also overexpressed. Our results provide important insights into the formation of fine ridges on epidermal cells in plants using *H. trionum* as a model.

## Introduction

The genus *Hibiscus* belongs to the Malvaceae family and is widely distributed in both temperate and tropical regions of the world. There are over 200 species in the genus, ranging from annual and perennial flowers to woody shrubs and small trees. *Hibiscus syriacus* (known as the Rose of Sharon) and *Hibiscus rosa-sinensis* (China rose) are two common ornamental plants with attractive flowers. As reviewed in Da-Costa-Rocha et al. (2014)^1^, *Hibiscus sabdariffa* (Roselle) is native to Africa and is used worldwide for its fibers, food, and medicinal properties. Another African species, *Hibiscus cannabinus* (kenaf), is also known for its fiber production.

*Hibiscus trionum* is an annual to short-lived perennial plant, widespread in tropical and temperate regions of Europe, Asia, and Africa. Commonly known as the ’Flower of an Hour’, its flowers are only open for a few hours in the morning. The center of the flower is deep purple with anthocyanin pigments, and the pistil is fused with many stamens (Fig. 1A). The adaxial surface of the purple region is striated with fine structures like ridges that show iridescence^2, 3^ (Fig.1C-e and 1D-a1,a2). As the flower opens, the epidermal cells in the purple region elongate to form the fine structure through a combination of mechanical stress and chemical changes to the cuticle^4, 5^. On the other hand, the outside of the petal is light yellow with flavonols. The adaxial epidermal cells in the light yellow region are conically shaped (Fig. 1C-bc), and the cells at the most distal tip are almost pointed (Fig. 1C-a). In contrast to the purple region, no ridges were observed in the light yellow region^2, 4, 5^ (Fig. 1C and 1D-d1,d2).

**Figure 1.**
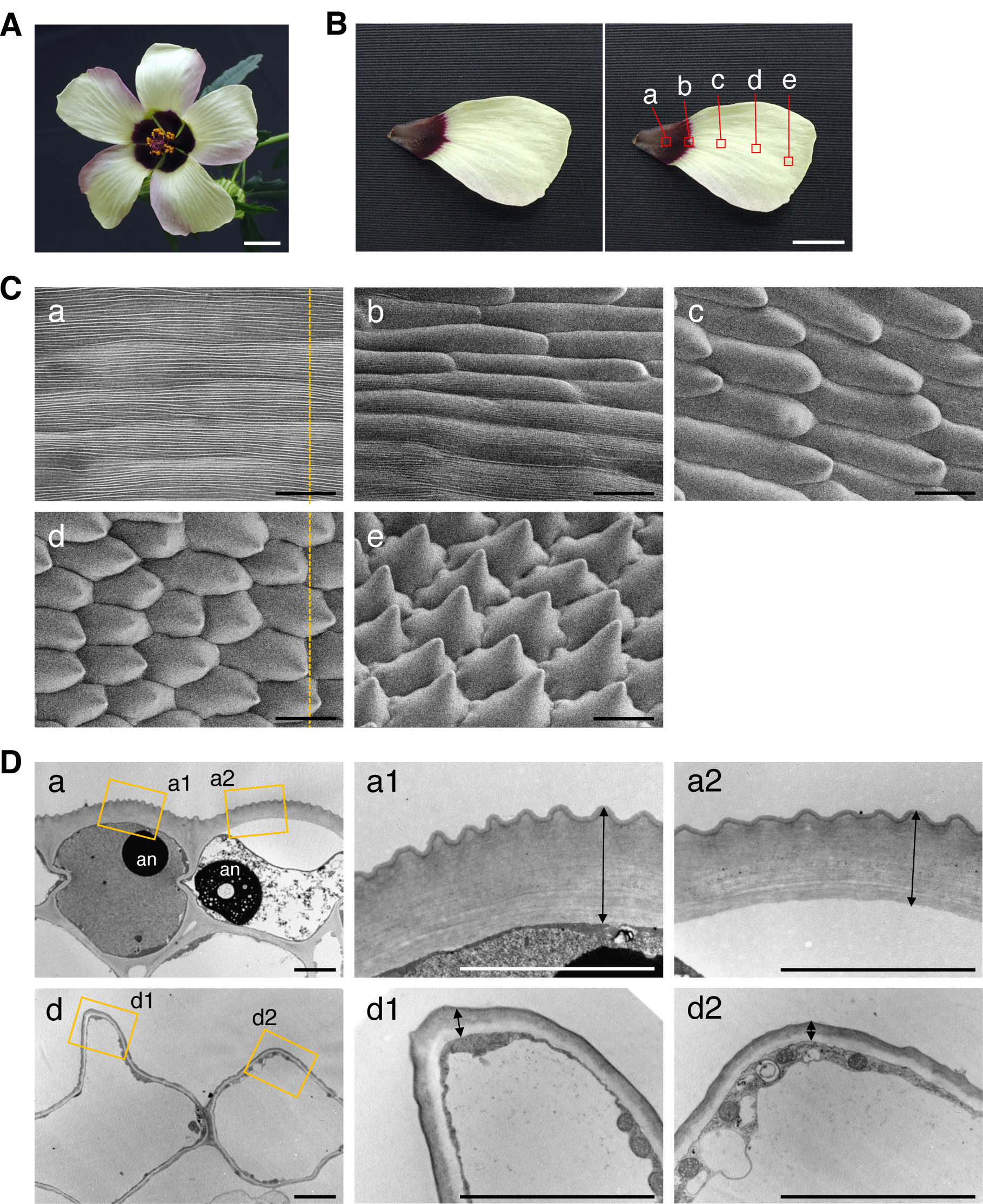
Flowers of *H. trionum* and their electron microscopy images. (A) A flower image of *H. trionum*. (B) A petal image of *H. trionum*. The right image shows the observation positions in C and D. (C) Scanning electron microscopy images at each position indicated in B. The yellow broken lines indicate the sectioning axis in D. (D) Transmission electron microscopy images obtained for positions a and d indicated in B, along the sectioning axis shown in C. a1 and a2 in a, and d1 and d2 in d are magnified views of the positions indicated by the yellow rectangles. Arrows in a1, a2, d1, and d2 indicate extracellular matrix including cell wall and cuticle layers. an, anthocyanin. Scale bars, 1cm in A,B; 50 μm in C,D.

The *H. trionum* genome provides a valuable resource for investigating the molecular mechanism of the formation of fine structures with iridescence, for supporting omics analyses (i.e. transcriptomics, proteomics, and metabolomics), and genome editing. In our incubator, *H. trionum* flowers when it reaches a plant length of about 15 cm, without the need for long or short day photoperiods. Furthermore, self-fertilization is possible with a lifecycle of about two months. Therefore, *H. trionum* has the potential to be a model plant for the study of flower iridescence.

In this study, we sequenced the genome of *H. trionum* using PacBio Sequel II systems powered with HiFi sequencing technology, which can produce highly accurate long reads. We also compared its transcriptomes between petal regions with and without the fine ridges to identify candidate genes. Our results provide important insights into the formation of fine structures on epidermal cells with iridescence.

## Materials and methods

### Plant materials and culture conditions

The seeds of *Hibiscus trionum* L. were obtained from the botanical gardens of Osaka Metropolitan University (Osaka, Japan), Tokyo Metropolitan Medicinal Plants Garden (Tokyo, Japan), Tokyo Metropolitan Kiba Park (Tokyo, Japan), Shigei Medicinal Plants Garden (Kurashiki, Japan), and a flower shop in Hamamatsu (Hamamatsu, Japan). Plants were cultivated at 22-25 °C under long-day conditions (16 h light and 8 h dark) in the soil.

### Preparation of chromosomes

Root tips were pretreated with 2 mM 8-hydroxyquinoline for 4 h at room temperature in the dark conditions. The pretreated root tips were fixed with 3:1 (v/v) ethanol/glacial acetic acid for 3 days at room temperature and stored at 4 °C until use. The fixed root tips (two to three) were then soaked in distilled water for 10 min and the water was removed with a Kimwipe on the glass slide. The root tips were stained with 2% (w/v) carmine (Merck, NJ, USA) in 45% acetic acid for 1 h at room temperature and washed with distilled water for 2 min to remove the acetic acid. Two to three root tips were put on the glass slides and the water was removed with a Kimwipe. Then, treated with enzymatic solutions consisting of 2% (w/v) of pectolyase Y23 (Kyowa Kasei, Osaka, Japan), 2% (w/v) of cytohelicase (Sigma-Aldrich, MO, USA), and 2% (w/v) of cellulase “ONOZUKA” R-10 (Yakult, Tokyo, Japan) dissolved in 1 x citrate buffer (0.01 M of Na-critic and citric acid diluted with distilled water and adjusted to pH 4.5-4.8) at 37 °C for 30 min to 1 h depending on the root size. Ten microliters of 45% acetic acid were added to the root tips and the acetic acid squash method was used with a 22 × 22-mm coverslip. Images were captured with a DP21 camera (Olympus/Evident, Tokyo, Japan) using a BX40 microscope (Olympus/Evident, Tokyo, Japan), and the number of chromosomes was counted manually.

### Flow cytometric analysis

For ploidy analysis, leaves cut into 5 mm squares were sectioned with a blade in 200 μl nuclear extraction buffer of the Quantum Stain NA UV2 kit (Quantum Analysis, Münster, Germany) containing 2 μl/ml 2-mercaptoethanol and 2% PVP K30, and incubated for 1 min at room temperature. 1 ml of DAPI staining solution from the same kit was added and filtered through a 30 μl nylon mesh. After incubation for 1 min at room temperature, the fluorescence intensity of DAPI was measured using a flow cytometer, Quantum P (Quantum Analysis, Münster, Germany).

For genome size estimation, leaves were subjected to the same treatment as for the ploidy analysis using the Quantum Stain PI kit (Quantum Analysis, Münster, Germany). 1 ml of PI staining solution from the same kit was added and filtered through a 30 μl nylon mesh. After incubation for 1 h in the dark conditions, the fluorescence intensity of PI was measured using a flow cytometer, CyFlow SL (Partec, Münster, Germany).

### Electron microscopy

For scanning electron microscopy, the raw samples of petals were observed using HITACHI Miniscope TM4000Plus (Hitachi High-Tech, Tokyo, Japan) with a cooling stage at −30 °C and an acceleration voltage of 5 kV.

For transmission electron microscopy, petals were fixed with 2% glutaraldehyde in 50 mM cacodylate buffer (pH 7.4) at 4°C overnight. The fixed samples were washed three times in 50 mM cacodylate buffer and then post-fixed with 2% osmium tetroxide in the same buffer for 3 h at 4°C. After dehydration in a graded ethanol series (50, 70, 90, and three times with 100%), the samples were infiltrated with propylene oxide (PO) for 1.5 h, a 50:50 mixture of PO and Epon812 resin (TAAB, Berks, England) for 3 h, and 100% Epon812 resin overnight, and polymerized at 60°C for 1 day. Ultrathin sections (70 nm thickness) were cut with a diamond knife (Diatome, Nidau, Switzerland) on a Leica EM UC6+FC6 ultramicrotome (Leica Microsystems, Wetzlar, Germany) and mounted on copper grids. The sections were stained with 2% uranyl acetate for 10 min followed by lead staining solution (Sigma-Aldrich, Massachusetts, USA) for 3 min. The grids were observed under a transmission electron microscope JEM-1010 (JEOL, Tokyo, Japan) with a CCD camera at an acceleration voltage of 80 kV.

### Isolation of high-molecular-weight genomic DNA

Young fresh leaf tissues were frozen in liquid nitrogen and homogenized into powders. The 0.7 g tissue powders were washed twice with a previously described method using a 45 ml sorbitol wash buffer [100 mM Tris-HCl pH 8.0, 0.35 M sorbitol, 5 mM EDTA pH 8.0, 1% (w/v) polyvinylpyrrolidone (PVP-40), and 1% (v/v) β-mercaptoethanol] to remove polysaccharides^6^. The IBTB buffer was prepared by mixing the isolation buffer [IB; 15 mM Tris, 10 mM EDTA, 130 mM KCI, 20 mM NaCl, and 8% (w/v) PVP-10, pH9.4], 0.1% Triton X-100, and 7.5% (v/v) β-mercaptoethanol. The obtained pellet of 1.4 g tissue was mixed vigorously with 14 ml of ice-cold IBTB buffer in a 50 ml conical tube. The mixture was filtered through a 100 μm nylon filter, followed by a 40 μm filter to remove tissue fragments. The filtrate was gently mixed with 140 μl of Triton X-100, and centrifuged at 4°C for 10 min at 2000 ×g to pellet the nuclei. The nuclear pellet from 2.8 g of tissue was vortexed with 10 ml of 2× CTAB buffer [2% (w/v) hexadecyltrimethylammonium bromide (CTAB), 1.4 M NaCl, 20 mM EDTA, 100 mM Tris-HCl (pH 8.0)] and 1 mg/ml Proteinase K, and incubated at 50°C for 30 min with gentle agitation. RNase A was added to the mixture to achieve a final concentration of 0.1 mg/ml, and incubated at 37°C for 30 min. After the addition of 10 ml of chloroform, the mixture was rotated for 30 min and centrifuged at 7,500×g for 15 min. The upper aqueous phase was transferred to a new tube, and the same procedure was repeated. The aqueous phase was combined with an equal volume of 2-propanol and centrifuged at 4,500×g for 20 min. The precipitate was rinsed twice with 3 ml of 70% ethanol, and air-dried for 10 min and then resuspended in 50 µl of TE [10 mM Tris-HCl (pH 8.0), 1 mM EDTA].

### Library preparation and sequencing

The genomic DNA library was constructed using the SMRTbell Express Template Prep kit v2.0 (PacBio, CA, USA) with a size range of 10-50kbp using the BluePippin size selection system (Sage Science, Massachusetts, USA). The libraries were sequenced on two cells of the PacBio Sequel II platform using the Sequel II Binding kit v2.2 (PacBio, CA, USA).

### Genome assembly and gene prediction

HiFi reads were assembled using Hifiasm^7^ (v0.16.1) with the ‘-N 1000’ option, then chloroplast sequences were removed. The assembled genome was evaluated using BUSCO^8^ (v5.4.4) with the eudicots_odb10 data set. Gene prediction was performed by BRAKER^9, 10^ using *H. trionum* RNA-seq data (the same datasets as in the transcriptome analysis section) and plant protein datasets obtained from OrthoDB^11^ (v11). The predicted genes obtained from each dataset were merged using TSEBRA^12^, and genes with at least partial support from known sequences were selected using a script from BRAKER, selectSupportedSubsets.py, with the ‘--anySupport’ option. The final version of the predicted genes was evaluated by BUSCO using the eudicots_odb10 data set.

### Gene annotation

The predicted gene sequences were translated to protein sequences using GffRead^13^ (v0.12.7). The protein sequences were BLASTP searched^14, 15^ (v2.13.0+) against *Arabidopsis thaliana* protein sequences. The results were filtered with the threshold (query coverage >= 80 and e-value <= 1e-5) and the descriptions of the best-hit proteins in *A. thaliana* were mapped to *H. trionum* sequences. Ortholog groups were predicted using OrthoFinder^16^ (v2.5.4) with the protein sequences of the species shown in Supplementary Table S1. Transcription factors (TFs) were searched based on the annotations of *A. thaliana* included in each ortholog group. The *A. thaliana* genes with the Gene Ontology term^17^ “DNA-binding transcription factor activity” (GO:0003700) were selected, and the gene members within the same ortholog groups were identified as TF-encoding genes. The classification of TF families was performed based on PANTHER annotations^18^ in *A. thaliana* and then manually curated.

### Transcriptome analysis

Petals of 5-6 mm in size were collected and dissected into the purple and light yellow regions. RNA was extracted using an ISOSPIN Plant RNA kit (Nippon Gene, Tokyo, Japan). The RNA integrity was checked using the Agilent 2100 Bioanalyzer System (Agilent Technologies, CA, USA) with an RNA Integrity Number (RIN) value greater than or equal to seven. Sequencing libraries were constructed using the TruSeq stranded mRNA library kit (Illumina, CA, USA) after random fragmentation of the reverse-transcribed cDNA. The libraries were sequenced on a NovaSeq 6000 system (Illumina, CA, USA). The obtained reads were evaluated using FastQC^19^ (v0.11.5), and adapter, poly-A, and low-quality sequences were removed using Cutadapt^20^ (v2.5). The remaining reads were mapped to the *H. trionum* genome with the gene structure annotations, and transcript per million (TPM) were calculated using RSEM^21^ (v1.3.1) with a mapping tool STAR^22^ (v2.7.10b). The differential expression genes (DEGs) between the purple and light yellow regions of the petals were identified using TCC^23^ of the R software.

### Pathway analysis

Pathways related to cuticle and anthocyanin were searched in *A. thaliana*, and the lists of genes involved in these pathways were obtained, using the Kyoto Encyclopedia of Genes and Genomes (KEGG) database^24^. The gene lists were then integrated with the *A. thaliana* annotations assigned to the *H. trionum* genes.

### Phylogenetic analysis

BLASTP searches were performed against the protein sequences of the species listed in Supplementary Table S1 using AtSHINE1, 2, and 3 as query sequences. The results were filtered with a threshold, e-value < 1e-50. Multiple sequence alignment was performed using MAFFT^25^ with the accurate option L-INS-i. A phylogenetic tree was constructed based on the multiple sequence alignment with complete deletion of gap sites using the maximum-likelihood method of MEGA 11 software^26^, with 1,000 bootstrap replicates. Jones-Taylor-Thornton (JTT) matrix-based models^27^ with gamma-distribution among sites and subtree-pruning-regrafting – extensive (SPR level 5) were used for amino acid substitution modeling and heuristic methods. The alignment file in FASTA format is provided as Supplementary Dataset S1.

## Results

### Selection of a diploid *Hibiscus trionum* line and its genome size

In *Hibiscus trionum*, both diploid (2n = 2x = 28) and tetraploid (2n = 4x = 56) lines have been reported^28–31^. For efficient genetic analyses, we focused on the diploid line. Five *H. trionum* lines were collected from the following locations in Japan: The botanical garden of Osaka Metropolitan University (Osaka line), Tokyo Metropolitan Medicinal Plants Garden (Tokyo-Medicinal line), Tokyo Metropolitan Kiba Park (Tokyo-Kiba line), Shigei Medicinal Plants Garden (Kurashiki line), and a flower shop in Hamamatsu (Hamamatsu line). Osaka line confirmed the tetraploid number (2n = 2x = 56) of the chromosome by chromosome counting (Supplementary Fig. S1A). We compared DAPI fluorescence intensities by flow cytometry with other lines with the tetraploid Osaka line to confirm the ploidy. The fluorescence intensity peaks of the Tokyo-Medicinal, Tokyo-Kiba, and Kurashiki lines were detected at the same position as the Osaka line, whereas the peak of the Hamamatsu line was at half the position (Supplementary Fig. S1B). Based on these results, the Hamamatsu line was considered to be diploid.

We estimated the genome size of the diploid line (Hamamatsu line) by comparing the peaks of PI fluorescence intensity with those of *A. thaliana* (135 Mb) and *Nicotiana benthamiana* (about 3 Gb) using flow cytometry. The relative peak positions were 67.33 and 209.13 between the Hamamatsu line and the *A. thaliana* (Supplementary Fig. S1C) and 210.43 and 344.94 between the Hamamatsu line and the *N. benthamiana*, respectively (Supplementary Fig. S1D). Based on these ratios, the genome size of the diploid *H. trionum* line was estimated to be 1.6-1.8 Gb.

### Genome assembly and the gene annotation

Extracted high-molecular-weight genomic DNA from the diploid line was sequenced using the Sequel II system (PacBio, CA, USA). The obtained HiFi reads were assembled using the Hifiasm assembler as shown in Table 1. The total length of the contigs was about 1.67 Gb, which was consistent with the estimated size. The N50 length and the number of contigs were about 41 kb and 276, respectively. The BUSCO evaluation showed 99.5% coverage of the core genes.

**Table 1.**
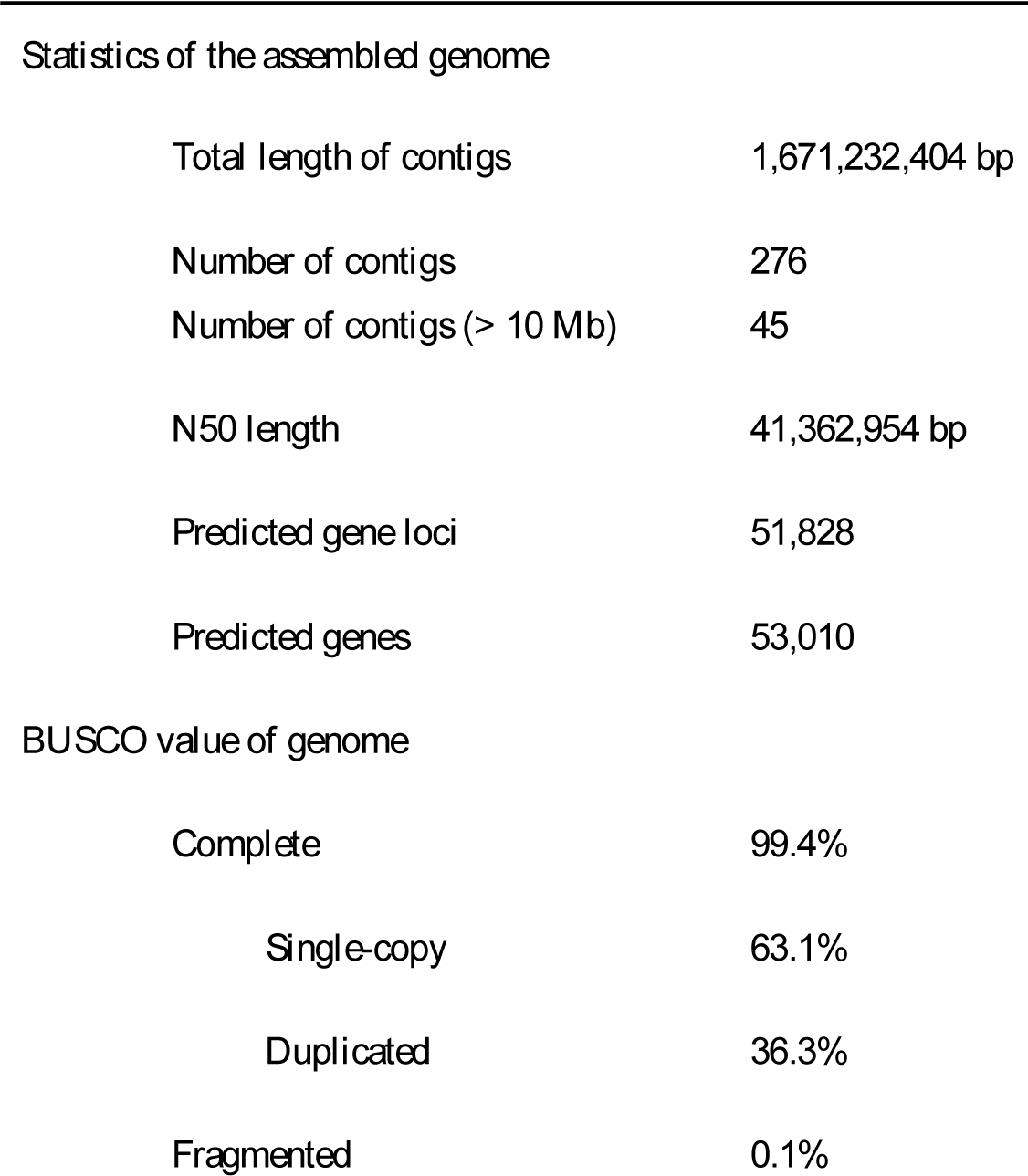
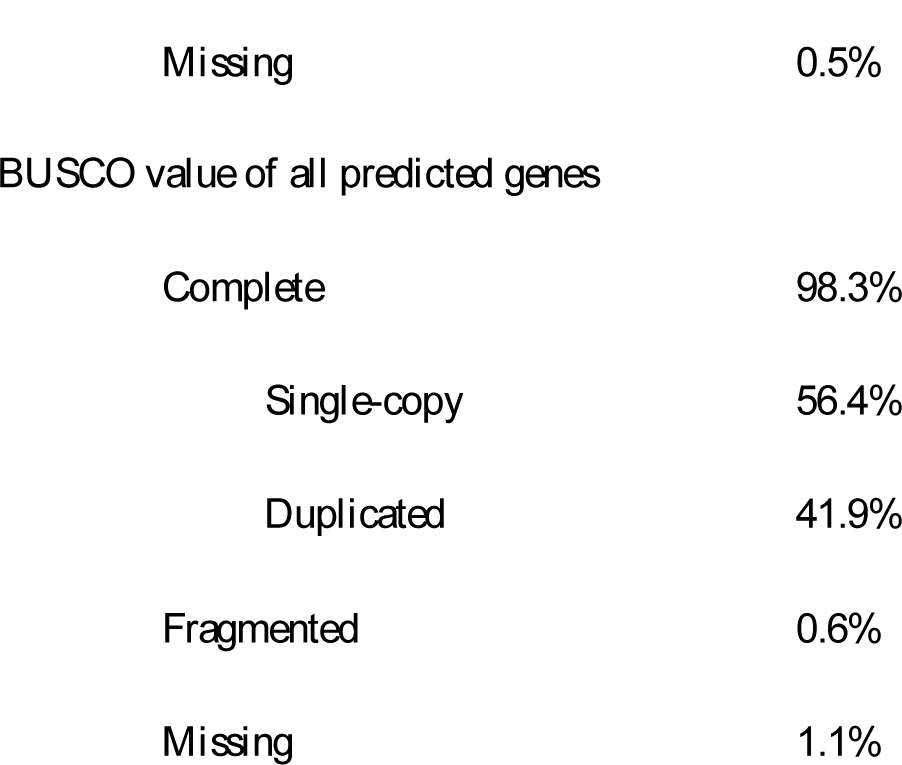
Statistics and BUSCO value of the assembled genome.

|  |  |
| --- | --- |
| Total length of contigs | 1,671,232,404 bp |
| Number of contigs | 276 |
| Number of contigs (> 10 Mb) | 45 |
| N50 length | 41,362,954 bp |
| Predicted gene loci | 51,828 |
| Predicted genes | 53,010 |
| BUSCO value of genome |  |
| Complete | 99.4% |
| Single-copy | 63.1% |
| Duplicated | 36.3% |
| Fragmented | 0.1% |
| Missing | 0.5% |
| BUSCO value of all predicted genes |  |
| Complete | 98.3% |
| Single-copy | 56.4% |
| Duplicated | 41.9% |
| Fragmented | 0.6% |
| Missing | 1.1% |

Gene prediction was performed by BRAKER using our RNA-seq data (the same data as in the next subsection) and the protein datasets from OrthoDB. In total, 51,828 loci and 53,010 genes were predicted with support from known sequences, and these genes included 98.3% of the BUSCO core genes (Table 1). The predicted genes were further annotated with BLASTP searches between the translated protein sequences and *A. thaliana* protein sequences.

Next, we examined the presence of major TFs. Ortholog groups were created among nine plant species (*H. trionum*, *Hibiscus syriacus*, *A. thaliana*, *Solanum lycopersicum*, *Ipomoea nil*, *Lotus japonicus*, *Antirrhinum majus*, *Populus trichocarpa*, and *Oryza sativa*) (Supplementary Table S2), and TFs were identified based on the annotation of *A. thaliana*. The resulting number of major TFs in *A. thaliana*, *H. trionum*, and *H. syriacus* is shown in Table 2. The details of all TF members can be found in Supplementary Table S3.

**Table 2.** The number of TFs in each species.

| TF family | *A. thaliana* | *H. trionum* | *H. syriacus* |
| --- | --- | --- | --- |
| MADS | 119 | 86 | 181 |
| MYB | 133 | 254 | 518 |
| NAC | 90 | 118 | 285 |
| WRKY | 68 | 159 | 323 |
| bHLH | 134 | 286 | 553 |
| bZIP | 51 | 104 | 221 |
| ARF | 23 | 51 | 97 |

### Comparison of gene expression between parts of petals with/without fine structures

To investigate the gene expressions in the purple region with the fine structures and the light yellow region without the fine structures, we performed RNA-seq analysis on these samples. Since the ridge structures are formed during flower development^4^, we collected petals before the completion of the fine structure (5-6 mm in size) and attempted to identify the responsible genes. The obtained Illumina short reads were mapped to the *H. trionum* genome sequence and differentially expressed genes (DEGs) were selected using the following threshold: q-value (false discovery rate [FDR]) < 0.001 and expression level ratio >= 2-fold. The highly expressed genes in the purple- and light yellow regions were 260 and 221, respectively (Supplementary Tables S4 and S5).

We characterize DEGs using three lists of genes: TF genes (blue flags), genes with almost no expression in the other region (average of transcript per million [TPM] < 0.15; yellow flags), and genes with extreme expression difference (> 100-fold; orange flags) (R-T columns in Supplementary Tables S4 and S5). Among DEGs that were highly expressed in the purple region, the TF list included 10 genes encoding bHLH family genes, bZIP family genes, ARF family proteins, regulation factors of flower development including MADS domain proteins, and an ERF/AP2 protein similar to AtSHINE1 for wax biosynthesis. The number of no-expression genes in the light yellow region was 14. They partially overlapped with the TF genes such as putative orthologs of *AtAPETALA1*, *AtBHLH32*, and *AtSHINE1*. The others were putative orthologs of *AtXTH30* (xyloglucan endotransglucosylase/hydrolase 30), *AtGA20ox* (involved in gibberellin biosynthesis), and genes involved in osmotic pressure. The high-difference list contained 17 genes, and nine of which overlapped with the no-expression genes. Remarkably, five genes showed expression ratios greater than 1000-fold: two were Arabidopsis CUTIN SYNTHASE2 (*AtCUS2*; involved in cutin synthesis) like genes and two were Arabidopsis dihydroflavonol 4-reductase (*AtDFR*; involved in leucoanthocyanidin synthesis) like genes.

The DEGs that were highly expressed in the light yellow region were as follows. The TF list contained 36 genes, including MYB domain proteins, bHLH family proteins, and a YABBY family protein involved in polarity specification of adaxial/abaxial axis. Notably, a *AtMYB16*-like gene was detected that regulates the epidermal conical cells in petals^32^. The list of no-expression in the purple region included 10 genes such as Arabidopsis ASYMMETRIC LEAVES 2-LIKE 1 (*AtASL1*; involved in proximal-distal patterning). The high expression list contained nine genes including the *AtYABBY5*- and *AtASL1*-like genes listed above.

### Pathway analyses of cuticle and flavonoid

We examined genes in the cuticle and flavonoid pathways that were expressed differentially between the purple and light yellow regions of *H. trionum* petals. We first extracted genes involved in the cuticle and flavonoid pathways in *A. thaliana*, and then located *H. trionum* genes based on the putative ortholog groups. *A. thaliana* genes orthologous to DEGs were highlighted in purple and yellow, respectively, based on their highly expressed region, respectively (Supplementary Tables S6 and S7).

In the cuticle pathway, we detected one *A. thaliana* gene orthologous to highly expressed genes in the light yellow region each in the fatty acid biosynthesis (ath00061) and in the fatty acid elongation (ath00062), and a few *A. thaliana* genes orthologous to highly expressed genes in both the purple and light yellow regions in fatty acid degradation (ath00071). We also found several *A. thaliana* genes orthologous to highly expressed genes in the purple and light yellow regions in cutin, suberine and wax biosynthesis (ath00073), but they were involved in the synthesis of the different metabolites. The genes highly expressed in the purple were putative orthologs of Arabidopsis cytochrome P450, family 77, subfamily A (*CYP77A* member) and HXXXD-type acyl-transferase family genes (*AtRWP1*), which are involved in the metabolism of cutin monomer, 10,16-dihydroxyhexadecanoic acid (10,16-DHPA) or polyhydroxy-fatty acid, and suberine monomer, 16-feruloyloxypalmitic acid, respectively (Fig. 2A). On the other hand, the gene highly expressed in the light yellow was a putative ortholog of Arabidopsis glucose-methanol-choline (GMC) oxidoreductase family gene (*AtHTH*), which is involved in the metabolism of suberine monomer α,ω-dicarboxylic acids (Fig. 2A).

**Figure 2.**
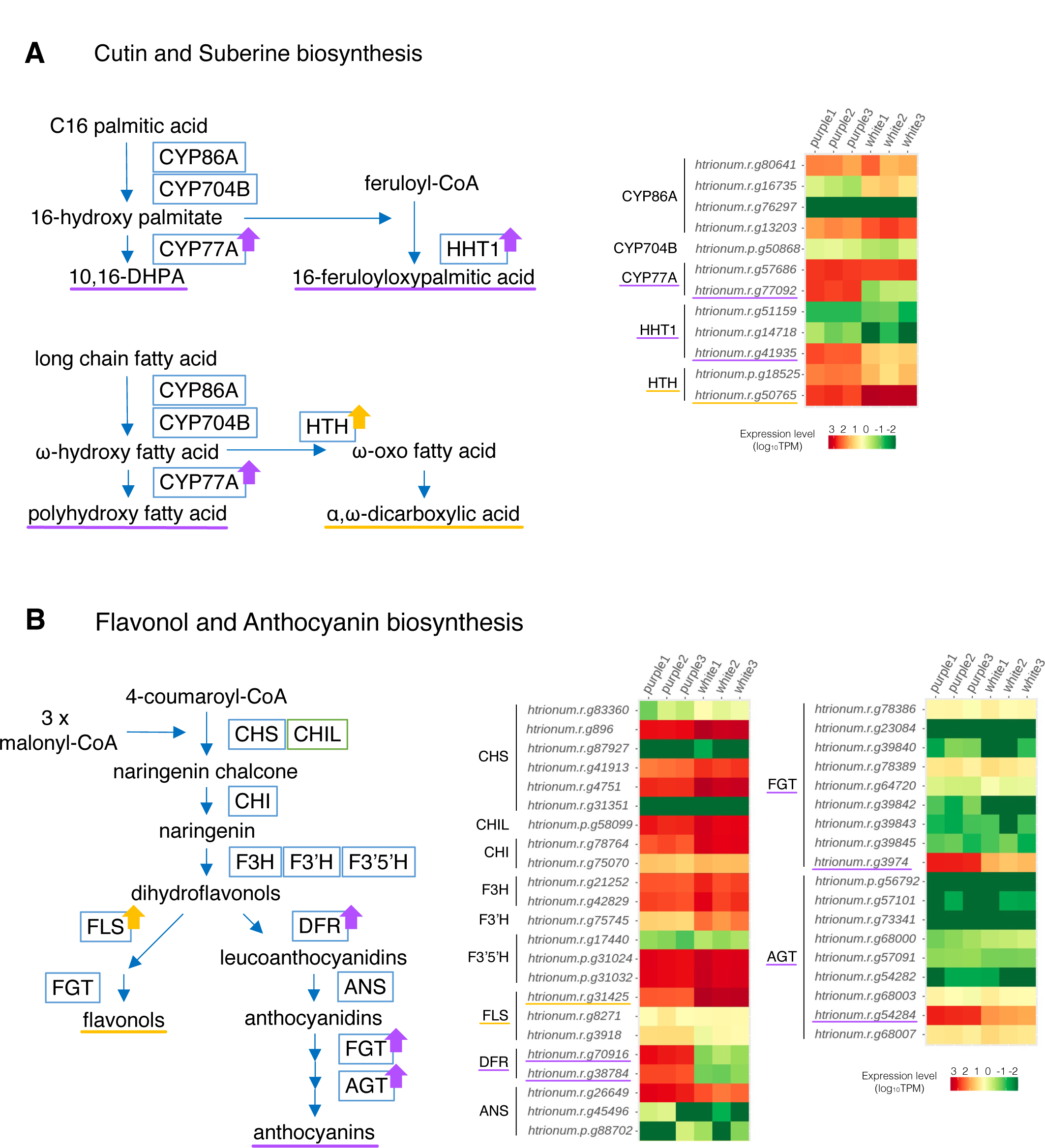
Biosynthesis pathways and expression profiles of enzyme genes. (A) Schematic diagram representation of the cutin and sberine biosynthesis (left), and the expression profiles of the enzyme genes (right). Enzymes are represented by blue rectangles. CYP86A, cytochrome P450 family 86 subfamily A; CYP704B, cytochrome P450 family 704 subfamily B; CYP77A, cytochrome P450 family 77 subfamily A; HHT1, omega-hydroxypalmitate O-feruloyl transferase; HTH, fatty acid omega-hydroxy dehydrogenase. (B) Schematic representation of flavonol and anthocyanidin biosynthesis (left), and the expression profiles of the enzyme genes (right). Enzymes and an enhancer are indicated by blue and green rectangles, respectively. CHS, chalcone synthase; CHI, chalcone isomerase; CHIL, chalcone isomerase-like protein, which is an enhancer of CHS^45, 46^; F3H, flavanone 3-hydroxylase; F3′H, flavonoid 3′-hydoroxylase; F3′5′H, flavonoid 3′,5′-hydoroxylase; FLS, flavonol synthase; DFR, dihydroflavonol 4-reductase; ANS, anthocyanidin synthase; FGT, fllavonoid glucosyl transferase; AGT, anthocyanin glucosyl transferase. The purple and yellow arrows in A and B indicate up-regulated enzymes in the purple and light yellow regions, respectively.

In the flavonoid pathway, we found one *A. thaliana* gene orthologous to highly expressed genes in the purple region each in the anthocyanin biosynthesis (ath00942) and in the flavone and flavonol biosynthesis (ath00944). In addition, one *A. thaliana* gene orthologous to the highly expressed genes in the purple and light yellow regions was found each in the flavonoid biosynthesis (ath00941). Genes highly expressed in the purple region were a putative ortholog of *AtUGT78D* member, *AtUF3GT*, and *AtDFR* encoding flavonoid glucosyltransferase (FGT), anthocyanin glucosyltransferase (AGT), and dihydroflavonol 4-reductase, respectively. A gene highly expressed in the light yellow region was a putative ortholog of Arabidopsis flavonol synthase 1 (*AtFLS1*) (Fig. 2B).

## Discussion

In this study, we sequenced the diploid *H. trionum* genome (2n = 28), whose genome size was consistent with the size estimated by flow cytometric analysis, and predicted a total of 53,010 genes on 51,828 loci in the genome. The genome and the predicted genes covered 99.4% and 98.3% of the core genes, respectively (Table 1), indicating the validity of our approach including the assembly and annotation. Several species in the genus *Hibiscus* have been proposed to have undergone whole genome duplication (WGD) independently^33^. The genomes of the previously analyzed *Hibiscus* species were longer than 1 Gb, and the number of chromosomes was larger than that of *H. trionum* (Table 3). The smaller number of chromosomes contributes to the analysis of the independent rounds of ploidy increase in the genus. Comparing the number of major TFs with *A. thaliana*, we found that MYB domain proteins, WRKY family proteins, bHLH family proteins, bZIP family proteins, and ARF family proteins were detected more than 2-fold in both *H. trionum* and *H. syriacus* (Table 2).

**Table 3.** Genome size and chromosome number in Hibiscus species.

| Species | Genome size (Gb) | Chromosome number | References for genomes |
| --- | --- | --- | --- |
| *H. trionum* (diploid) | 1.6 | 2n = 28 | This study |
| *H. syriacus* | 1.75 | 2n = 80 | Kim et al., 2017 |
| *H. mutabilis* | 2.68 | 2n = 92 | Yang et al., 2022 |
| *H. cannabinus* | 1.1 | 2n = 36 | Zhang et al., 2020 |
| *H. hamabo* | 1.7 | 2n = 92 | Wang et al., 2022 |

Several key genes in flavonol and anthocyanin biosynthesis were detected from DEGs. The genes *htrionum.r.g38784* and *htrionum.r.g70916* were specifically expressed in the purple region, while *htrionum.r.g31425* was promoted only in the light yellow region. The former two genes and the latter gene were putative orthologs of *AtDFR* and *AtFLS1*, which are responsible for the synthesis of leucoanthocyanidin (leading to anthocyanin biosynthesis) and flavonols from dihydroflavonol, respectively (Fig. 2B). The genes *htrionum.r.g3974* and *htrionum.r.g54284*, which show increased expression in the purple region, were assigned to ortholog groups of *AtUGT78D*s and *AtUF3GT*, respectively, both involved in the subsequent anthocyanin biosynthesis. The former, *htrionum.r.g3974*, showed the highest similarity to *AtUGT78D2* (one of the FGTs), and its mutant showed reduced anthocyanin content^34^. The latter, *htrionum.r.g54284*, showed the closest similarity to *AtUGT79B3* (one of the AGTs), which is known to increase anthocyanin accumulation through its overexpression^35^. In addition, *htrionum.r.g54284* was also predicted to be orthologous to the *I. nil* gene *INIL03g17665* (*Dusky*). *Dusky* encodes anthocyanidin 3-O-glucoside-2’’-O-glucosyltransferase (3GGT), and its mutant showed reddish-brown flowers with reduced purple coloration^36^. These findings suggest that the accumulation of anthocyanins and flavonols in the purple and light yellow regions is regulated by the expression of the above genes, contributing to the colors observed in each petal region (Fig. 2B). Note that *htrionum.r.g3974* may be involved in flavonol biosynthesis, as it also shows similarity to *AtUGT78D1*, which encodes an FGT that uses flavonols as substrates^37^.

Regarding cuticle components, a total of 10 genes were detected by OrthoFinder as the *SHINE* clade of AP2 domain transcription factors. In *Arabidopsis*, *AtSHINE1*, *AtSHINE2*, and *AtSHINE3*, are known to promote a brilliant or shiny green leaf surface with increased cuticular wax^38^, and Moyroud et al. (2022)^5^ focused on this clade as the candidate factors for fine ridge formation in *H. trionum*. In addition to the several genes reported by Moyroud et al. (2022)^5^, we newly detected two putative orthologs of *AtSHINE1, htrionum.r.g51082* and *htrionum.r.g73448* with a high bootstrap support in the phylogenetic analysis (Supplementary Fig. S2). Among the 10 candidates, only *htrionum.r.g51082* was highly expressed in the purple region (Supplementary Table S8) and is a promising candidate for fine ridge formation.

The *AtCUS2*-like genes, *htrionum.r.g2803* and *htrionum.r.g8620*, were highly up-regulated in the purple region but had almost no expression in the light yellow region, with expression ratios exceeding 1000-fold. *AtCUS2* is known to play a crucial role in the development of ridge structures during sepal growth^39^. We also observed a drastic difference in the thickness of the extracellular matrix between the two regions (Fig. 1D). Although the cell wall and cuticle layers were unclear by transmission electron microscopy, the amount of cutin components in the cuticle layer likely reflects the results of the intense expression of these genes to form ridge structures. The pathway analysis revealed that the synthesis of 10,16-DHPA, polyhydroxy-fatty acid, and 16-feruloyloxypalmitic acid was upregulated in the purple region (Fig. 2A). Mazurek et al. (2017)^40^ reported a clear correlation between the amount of 10,16-DHPA and the cuticular ridges on the conical cells of Arabidopsis petals, suggesting that the *AtCYP77A*-like gene, *htrionum.r.g77092*, is also responsible for the ridges. In contrast, the synthesis of ω-oxo fatty acid was upregulated in the light yellow region (Fig. 2A), suggesting that the cutin and suberin composition in the cuticle may differ significantly between the purple and light yellow regions. This difference may contribute to the absence/presence of fine structures on the petal surface.

As mentioned above, mechanical stress associated with cell elongation has been reported to be important for the formation of fine structures^4, 5^. In *H. trionum* petals, cells in the purple regions were elongated in the proximal-distal direction, while those in the light yellow regions showed conical shapes^2, 3^. In the purple regions, *AtXTH30*-like (*htrionum.r.g22645*) and *AtGA20ox*-like (*htrionum.r.g53103*) genes, which are involved in tissue development and elongation^41–44^, were upregulated. Additionally, the *AtMYB16*-like gene (*htrionum.r.g50021*), which regulates epidermal conical cells in petals^32^, was upregulated in the light yellow region. The change in cell shape in each region may affect the formation of fine structures. Genes involved in axis determination were also upregulated in the light yellow region.

We performed genome, transcriptome, and pathway analyses and we successfully identified promising candidate genes involved in the formation of the fine structures on the epidermal cells in *H. trionum* petals. This plant grows easily in a laboratory and has a lifecycle of only about two months. With its genome sequence, *H. trionum* can be a useful model plant for molecular analysis of ridge formation. Analyzing the molecular functions of the identified genes and elucidating the mechanisms of fine structure formation are our next research tasks.

## Data availability

The genome sequence, the genome annotations, and the raw sequence reads were deposited at DDBJ/ENA/GenBank under accession number fPVP.

## Acknowledgments

We thank K. Yoshimoto (Meiji Univ.) for experimental support, K. Fukushima (Würzburg Univ.) for providing the genomic DNA isolation method, and T. Takeuchi, A. Kawada, K. Kuzunishi, and C. Takagi at NIBB for plant culture. We also thank the botanical gardens of Osaka Metropolitan University (Osaka, Japan), Tokyo Metropolitan Medicinal Plants Garden (Tokyo, Japan), Tokyo Metropolitan Kiba Park (Tokyo, Japan), and Shigei Medicinal Plants Garden (Kurashiki, Japan) for providing *H. trionum* seeds. Computations were partially performed on the NIG supercomputer at the ROIS National Institute of Genetics and the Data Integration and Analysis Facility at the National Institute for Basic Biology. Plant culture and SEM observation were supported by the Model Plant Section of Model Organisms Facility and Optics and Imaging Facility in NIBB Trans-Scale Biology Center. This work was supported by JSPS KAKENHI grants to KS (20H05909), JST ACT-X grant to SK (JPMJAX21B6), a grant from Sapporo Bioscience Foundation to SK, and NIBB Collaborative Research Program to SK (20-341, 21-223, 22NIBB308, and 23NIBB314).

## Author contributions

S.K. and M.A. conceived and designed the research in general. S.K. and A.H. performed and supervised the experiments. T.I. performed the chromosome observation. S.M., A.S., and K.S. performed the sequencing and assembly of the genome. S.K. and M.A. performed and supervised the computational analyses. All authors provide comments on the manuscript.

## Notes

### Competing Interest Statement

The authors have declared no competing interest.

