## Supplementary Fig. for "Genome and transcriptome analyses reveal genes involved in the formation of fine ridges on petal epidermal cells in *Hibiscus trionum*"

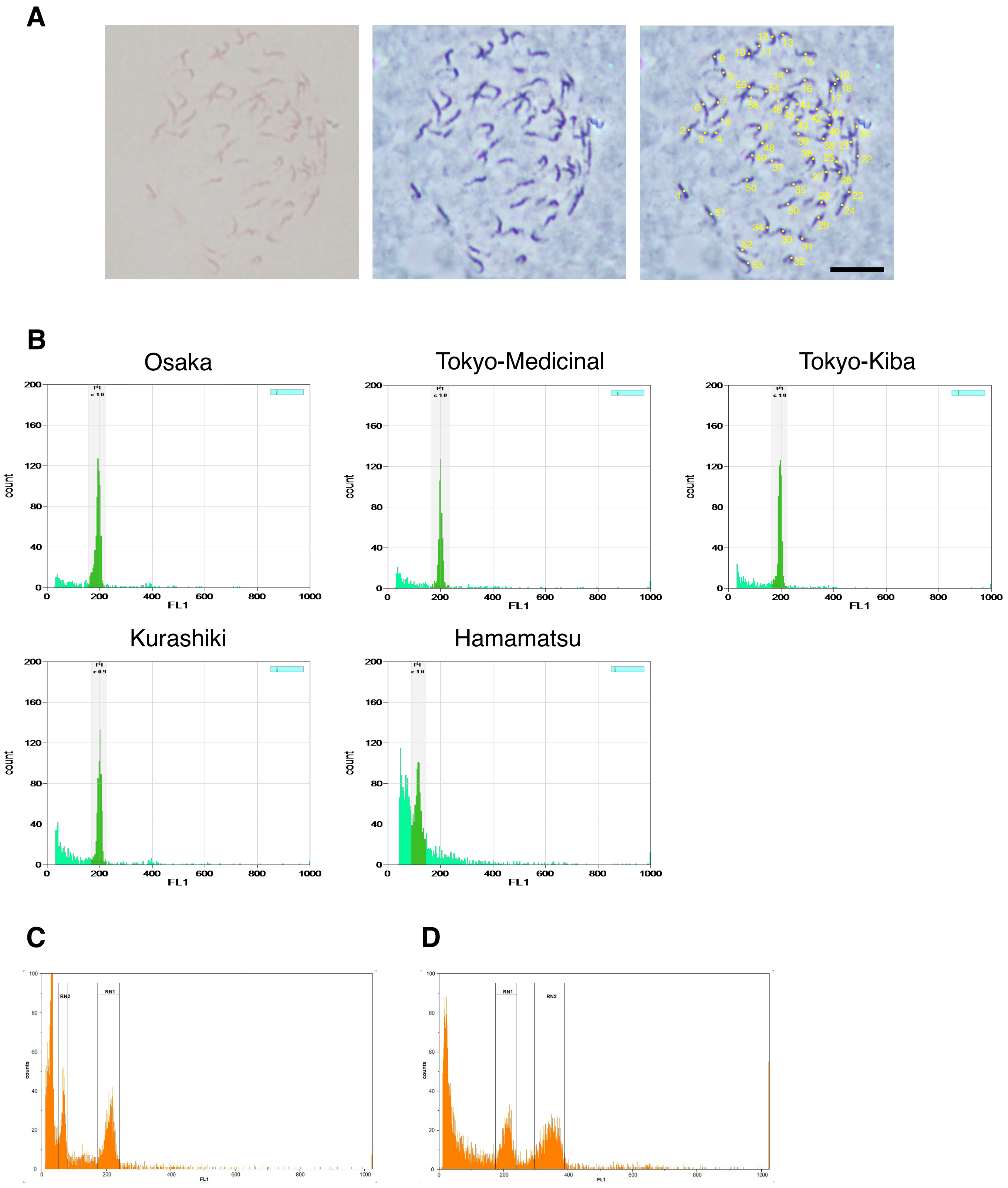


**Supplementary Figure S1.** Ploidy and genome size estimation in *H. trionum*. (A) Karyotype analysis in prophase using the Osaka line of *H. trionum*, showing 56 chromosomes. Yellow dots with numbers represent individual chromosomes. Scale bar = 10 µm. (B) Ploidy analysis with DAPI fluorescence signals by flow cytometry in each *H. trionum* line. Areas with gray backgrounds in each histogram indicate the peak positions. (C, D) Genome size estimation of the diploid *H. trionum* line (Hamamatsu line) through flow cytometry. RN1 with a peak of 67.33 and RN2 with a peak of 209.13 in C are Hamamatsu line and *A. thaliana* (8C = 540 Mb), respectively. RN1 with a peak of 210.43 and RN2 with a peak of 344.94 in D are Hamamatsu line and *N. benthamiana* (about 3 Gb).

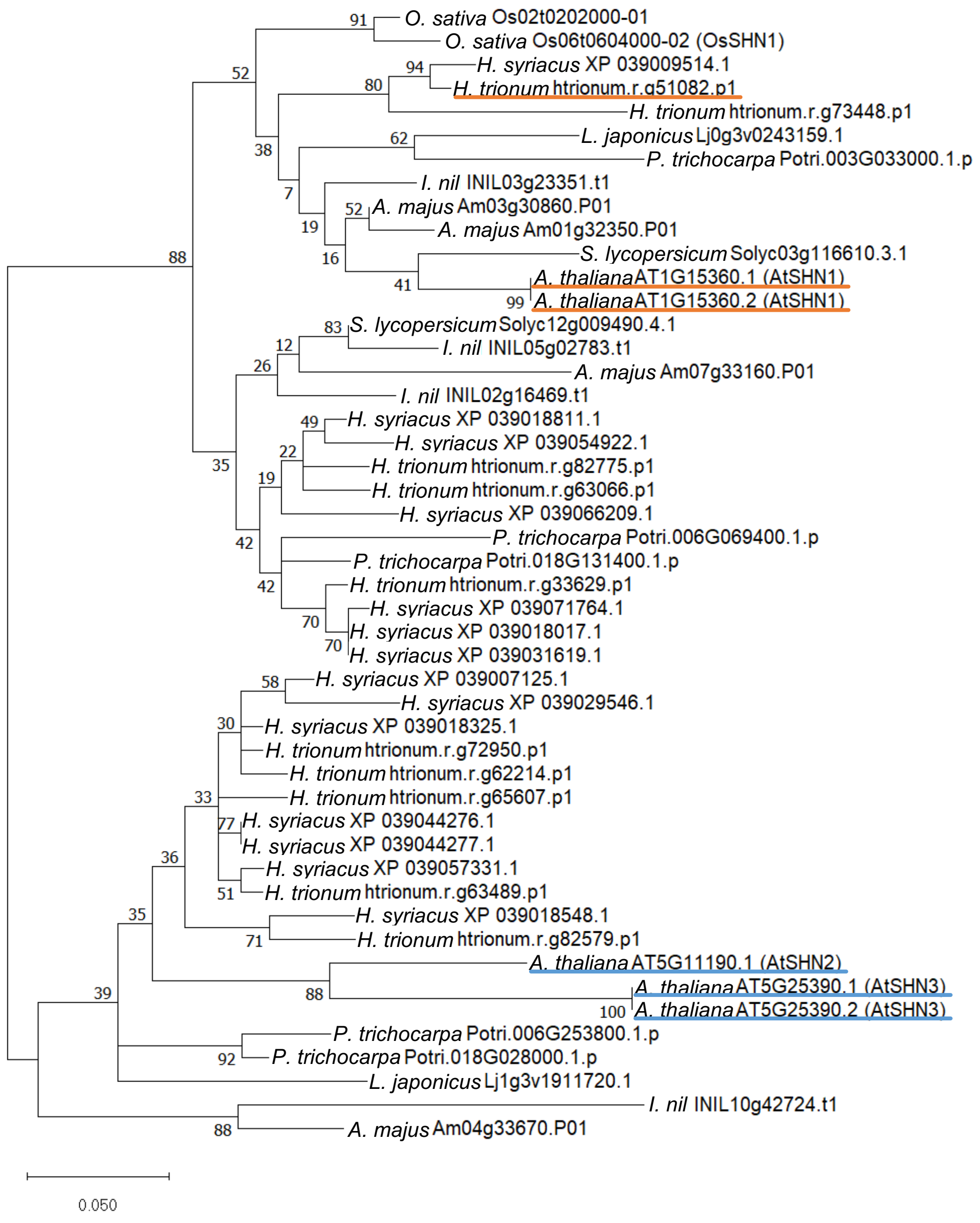


**Supplementary Figure S2.** Phylogenetic tree of the SHINE clade proteins. Branch lengths are proportional to the estimated number of amino acid substitutions per site (scale at lower left). The values shown at the nodes of the branches are bootstrap support values.
